# Assessment of Adherence to Antiretroviral Therapy Among Adult People Living with HIV/AIDS in North East, Ethiopia

**DOI:** 10.1101/492330

**Authors:** Almaz Mengistie, Adugnaw Birhane, Esubalew Tesfahun

## Abstract

**Back ground:** The scale-up of anti retroviral treatment is among the greatest successes of the global Acquired Immunodeficiency Syndromes (AIDS) response to date and it is on a Fast-Track- approach. To contribute to the global and local target of HIV /AIDS response as well as to obtain full benefits of ART medication, strong adherence of ≥95% to Anti-Retroviral Therapy (ART) is needed.

**Methods:** A cross-sectional study among 352 and 20 study participants for quantitative and for in-depth interview respectively. Data was collected using semi-structured questionnaire with a face-to-face interview for quantitative and interview guide for the in-depth interview. Bivariate and Multivariable logistic analysis was applied to examine the association between the dependent and independent variables. Additionally, thematic analysis, interpretation and triangulation of the finding were done for in-depth-interview.

**Results:** A total of 352 people living with HIV/AIDS were responded to the study with 90% response rate. Of the total 352 respondents 87.2 % were adherent and the overall adherence level was found to be 95%. The study revealed factors associated to adherence to ART were marital status(AOR:4.4;95%CI:1.3-14.7), use of memory aids (AOR:6.5;95%CI:2.6-16.2) Living condition (AOR: 2.9;95%CI: 1.1-7.6), Experienced side effect (AOR: 4.6;95%CI;1.9-11.3),drug regimen (AOR: 5.5; 95%CI: 1.4-22.3) and distance in km (AOR:2.7;95% CI; 1.11-6.4).

**Conclusion:** Patient counselling, health education and health system strengthening are very important for improvement of ART drug adherence.

## Introduction

The scale-up of anti retroviral treatment is among the greatest successes of the global Acquired Immunodeficiency Syndromes (AIDS) response to date and it is on a Fast-Track-Strategy. Its goal is to achieve and sustain viral suppression. Even though its roll-out has led to a massive reduction in AIDS-related deaths, viral suppression rates are still too low to realize the prevention role of treatment. The early initiation of ART may place demands on the health system in some settings that could increase the risk of drug resistance, drug stock-outs, insufficient patient preparation and suboptimal adherence. Adherence to an ARV treatment regimen involves taking all pills in the correctly prescribed doses, at the right time, and in the right way. To achieve the global and local target of HIV/AIDS response, strong adherence to Anti-Retroviral Therapy (ART) is needed to obtain full benefits of ART medication (1-3).

Adherence to ART is a primary determinant of viral suppression and transmission risk, disease progression and death. Long-term adherence to treatment is critical for the success of ART and presents new challenges. Suboptimal adherence is a major challenge in all regions, at all stages of HIV disease, and is associated with a diversity of patient- and program-related challenges. Study has suggested that when adherence rates are between 50% and 85%, drug resistance is more likely to develop (2-5).

The evidence in different countries and settings has shown sub optimal level of adherence. Non-adherence to ART is the most common reason for treatment failure and call for additional assessment and intervention.

Consequences of poor adherence to long-term therapies are poor health outcomes and increased health care costs. It severely compromises the effectiveness of treatment making this a critical issue in public health both from the perspective of quality of life and of health economics (6). It is the most common cause of treatment failure. Due to cross-resistance, the virus can become resistant to an entire class of ARVs thereby rendering that class ineffective not just for the individual but also for the society. This can lead to change of treatment regimen to more expensive second-line drug (7-9). Adherence is increasingly understood as a dynamic behaviour influenced by a matrix of interrelated factors that change over time and periodic assessment is recommended. Additionally, the rapid scale up and current early initiation of ART may be associated with an increase in HIV drug resistance in the population, if appropriate assessment and prevention measures are not taken. Because this early initiation leads people to start before they are ready, with adverse consequences for adherence and treatment outcomes. Because of these, evaluation of adherence was recommended in the literature in order to develop appropriate adherence interventions (2).

In order to implement appropriate intervention aimed to improve adherence, the discovery of factors for non-adherence to ART could help guide new public health policies and aid in developing more effective prevention strategies. In spite of the high non-adherence rate, there is very little information on the factors for these non-adherences. Thus, this study was conducted with an aim to assess the recent level of adherence to ART and investigate determinant factors. This study was conducted with an aim to assess the recent level of adherence to ART and investigate factors associated with adherence to ART medication using mixed method approach. Therefore, the result of this research project will have a paramount importance for the improvement of ART drug adherence intervention.

Therefore, the result of this research project will have a paramount importance for the improvement of ART drug adherence intervention.

## Methods

### Study Area and period

The study was conducted from February to April, 2018 at ART units in Debre Berhan Kebele Health Centre and Debre Berhan Referral Hospital. These two health institutions are the only public health facilities in the town currently giving ART service. According to the data from the two health facilities, there were a total of 2,644 adult patients actively following ART therapy in both health facilities (1,864 at the hospital and 780 at the health centre).

### Study Design and Data Collection

An institutional based cross-sectional quantitative study supported by qualitative approach was conducted. The data was collected using a standardized semi-structured questionnaire adopted from the World Health Organization (WHO) by directly interviewing study participants and gathering relevant information from their records. In addition, unstructured one-on-one in-depth interviews using interview guide were conducted at the study facilities with ART patients and healthcare workers and case managers administering ART services. The method used to measure adherence level was patient self-report on the number of doses skipped or missed on the past four days recall. The sample sizes for this study are 276 and 115 from the hospital and health centre respectively.

### Analysis

Self-reported adherence was classified as being “adherent” when not even a single dose is missed corresponds to dose adherence. If the patient admitted having missed at least one dose corresponds to non-adherence. Adherence measurement of 95% or more is classified as adherent and less than 95% is classified as having non-adherence.

Therefore, self-report of four-day recall adherence rates measured as proportion using the formula:

Adherence rate =Number of pills (doses) taken ÷ (doses prescribed or supposed to be taken × 100 for each respondents.

Descriptive statistics such as percentages, means, and standard deviations were calculated to describe and the data was also summarized in tables and graphs. To examine the relationship between levels of adherence and the independent variables, the bivariate and multivariable logistic regression analysis was carried out. The outcome ART adherence variable is dichotomized and defined as 0= none adherents to ART: 1= adherents to ART. Each independent variable was tested with the adherence status for the association. Thematic analysis was used for qualitative data. Interpreted and triangulation of the finding with quantitative finding was done and presented in text.

## Results

### Socio demographic characteristics

A total of 352 adult patients living with HIV/AIDS on first line ART were included in the study with the response rate of 90%. More than two third 249 (70.7%) of the respondents were females and the mean age of the respondents was 36.9 ± 10.6 years.

### Information regarding psycho social conditions

When the respondents asked about their living condition 242(68.8%) of them were living with their families (spouse or children). Regarding their disclosure status, 307(87.2%) were disclosed their sero status to their families, but only 134(38.1%) of the total respondents were disclosed their HIV status to the communities. About half 85(52.6%) of respondents reported they have discomfort of taking medication in front of others.

### Information regarding the clinical condition of the patient

About half of the patients 166(51%) had base line CD4 count less than 200 but; 210 (68%) of the participants had current CD4 count greater than 350. The initial WHO clinical stage was stage III for 131(37.3%) patients and the current WHO stage, for more than three fourth 336(95.5%)) of the respondents one was T1 (Table 1).

**Table 1:**
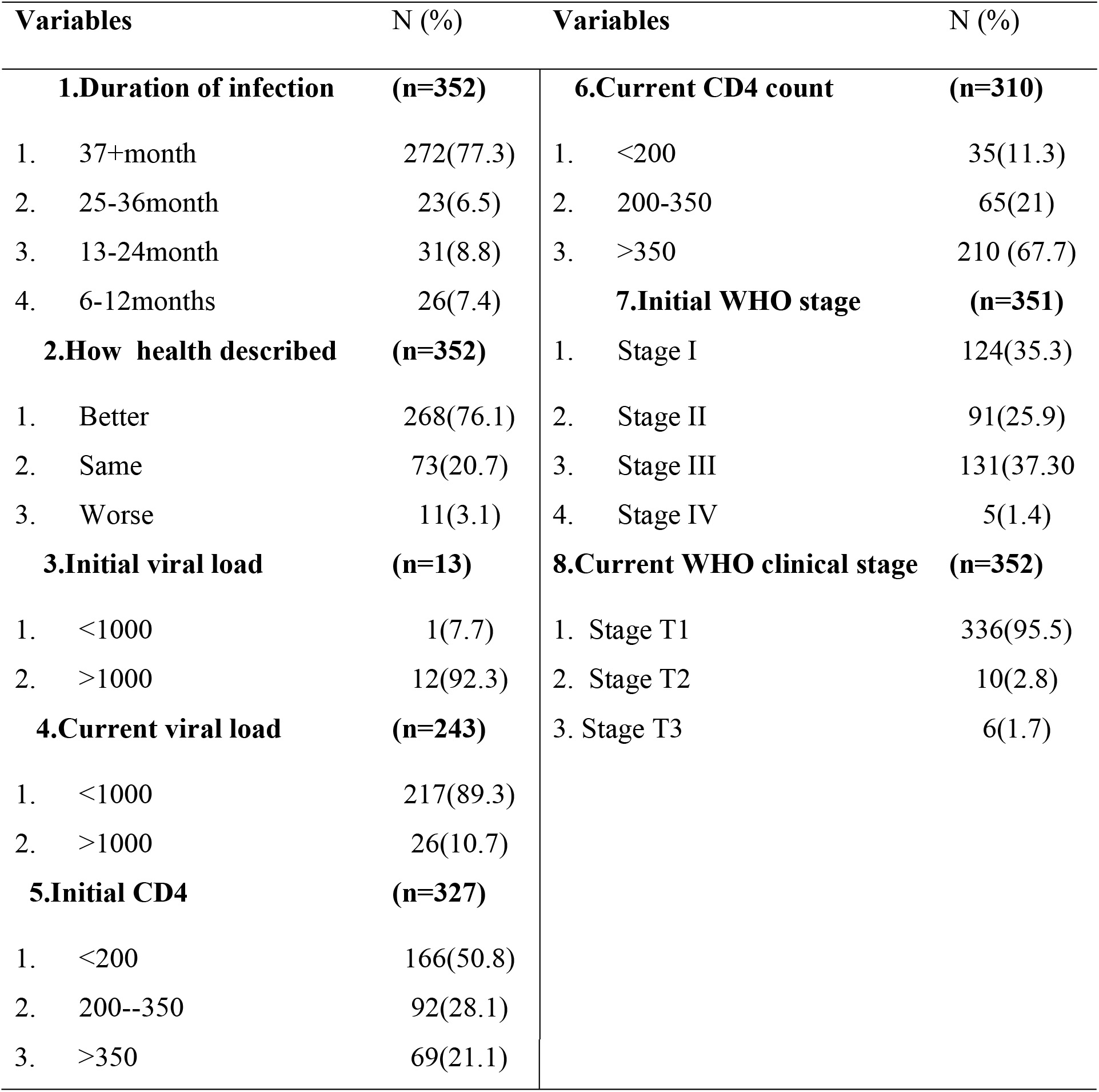
The clinical condition and clinical marker of respondents in Debre Berhan Health Centre and Debrebrahan Referral Hospital, North East Ethiopia, 2018

### Drug related

Of respondents on medication 194(55%) were taking one pill of ART once daily and the rest were taking twice during the study period. About 273 (77.9%) of them reported that the schedule was convenient and easy to fit to their daily routine, but the rests found it was inconvenient and difficult to fit to their daily routine (Table, 2).

**Table 2:**
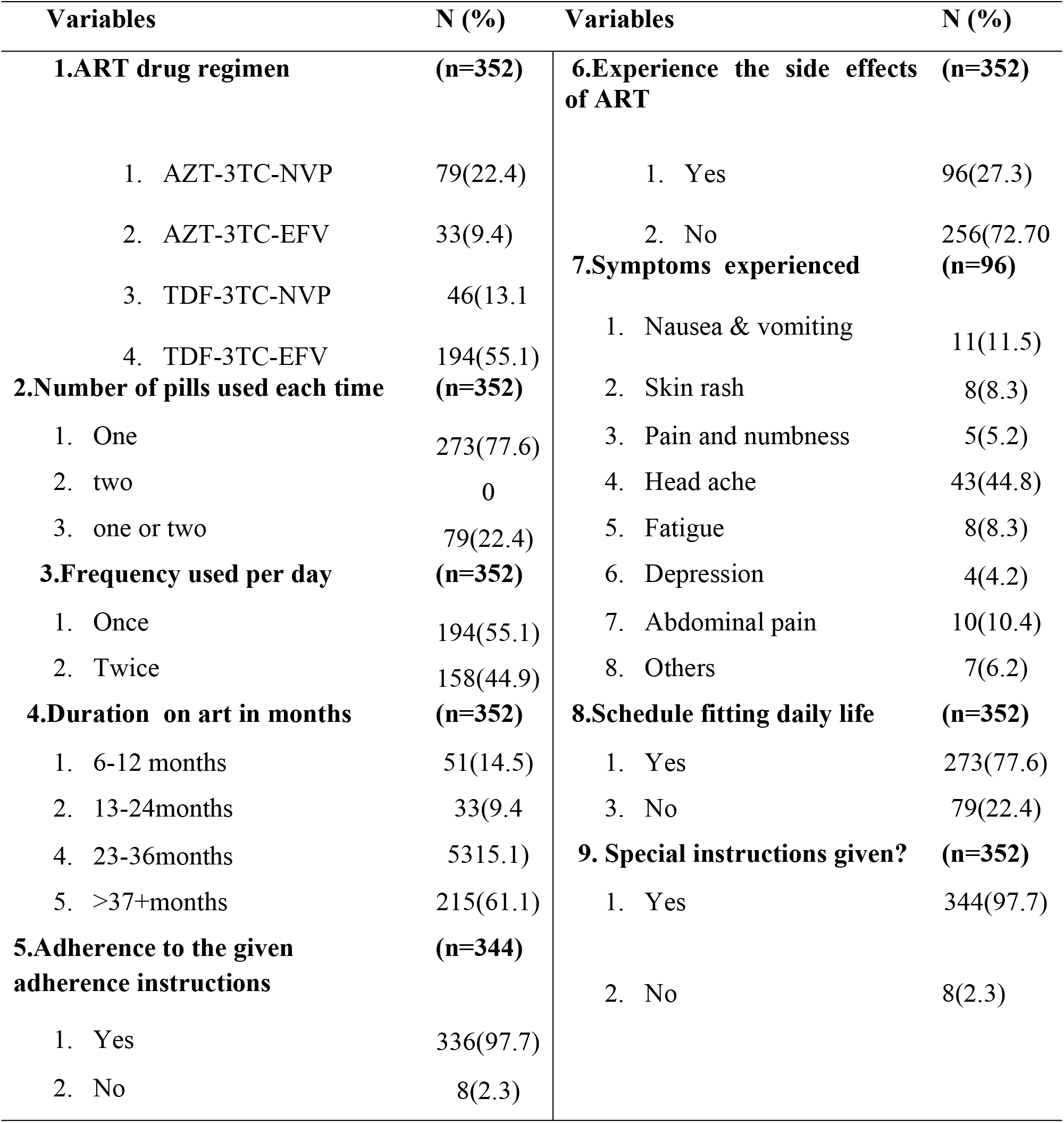
Drug related information of respondents in Debrebrahan Health Centre and Debrebrahan Referral Hospital, North East Ethiopia, 2018

### ART adherence level and reasons for missing their treatment

Regarding the adherence assessment to ARV therapy using missed dose self report medication adherence measurement, 307(87.2%) of all participants interviewed were adherent to the dose, and the rests 45(12.8%) were missed at least one dose in the last four days prior to the survey. The overall mean adherent level was 95%±14 in this study area. Of non adherents, 37(82.2%) of the study participant missed two doses (Table3). The primary reasons reported by participants for treatment non-adherence were 15(33.3%) due to simply forgotten, 12(26.7%) being away from home, being busy 7(15. 6%).

**Table 3:**
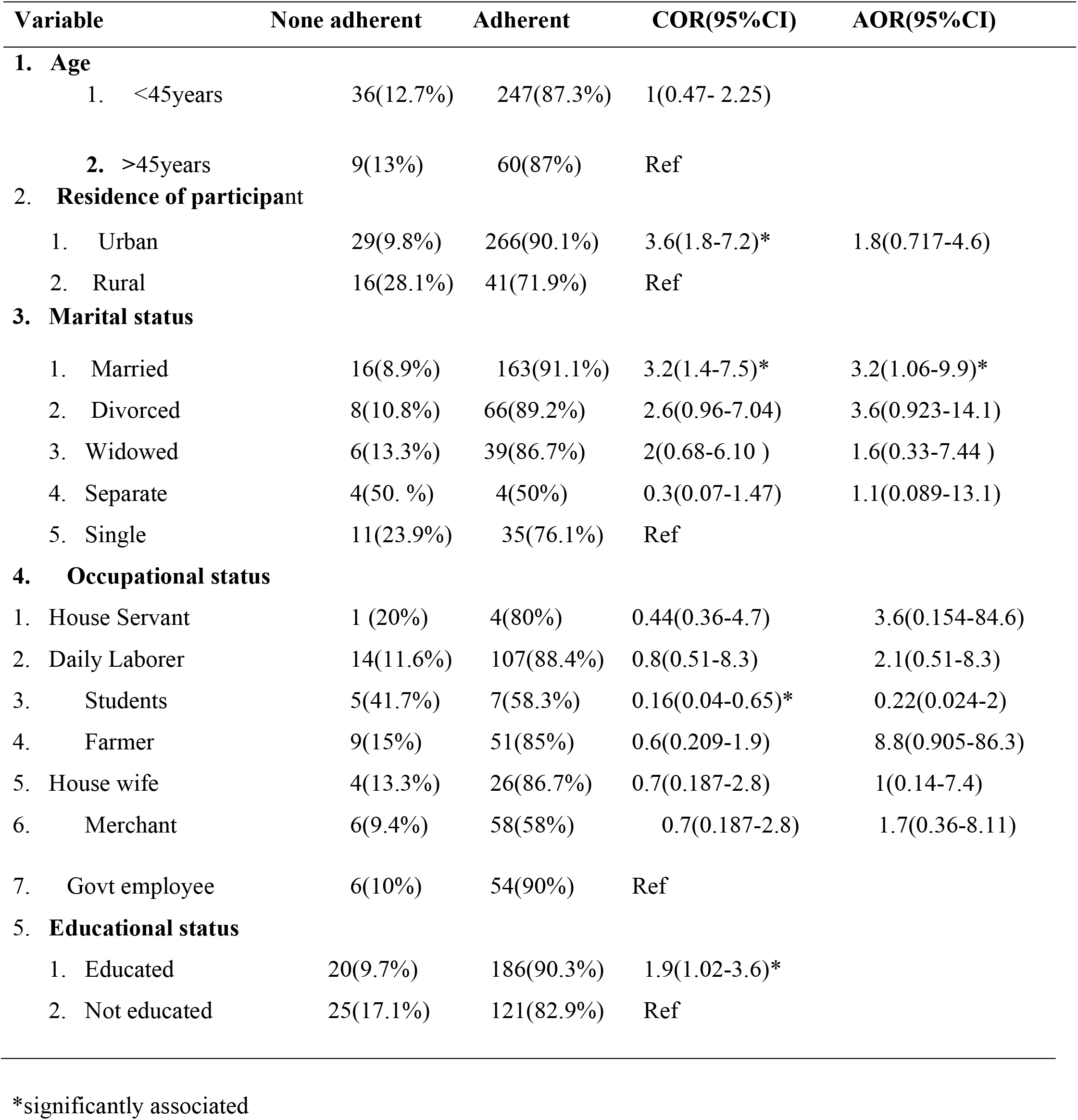
Bivariate and multivariate analysis of adherence and socio demographic variables of respondents in Debrebrahan Health Centre and Debrebrahan Referral Hospital, North East Ethiopia, 2018

| Variable | None adherent | Adherent | COR(95%CI) | AOR(95%CI) |
| --- | --- | --- | --- | --- |
| **1. Age** |  |  |  |  |
| 1. <45years | 36(12.7%) | 247(87.3%) | 1(0.47- 2.25) |  |
| 2. >45years | 9(13%) | 60(87%) | Ref |  |
| **2. Residence of participant** |  |  |  |  |
| 1. Urban | 29(9.8%) | 266(90.1%) | 3.6(1.8-7.2)* | 1.8(0.717-4.6) |
| 2. Rural | 16(28.1%) | 41(71.9%) | Ref |  |
| 3. Marital status |  |  |  |  |
| 1. Married | 16(8.9%) | 163(91.1%) | 3.2(1.4-7.5)* | 3.2(1.06-9.9)* |
| 2. Divorced | 8(10.8%) | 66(89.2%) | 2.6(0.96-7.04) | 3.6(0.923-14.1) |
| 3. Widowed | 6(13.3%) | 39(86.7%) | 2(0.68-6.10 ) | 1.6(0.33-7.44 ) |
| 4. Separate | 4(50. %) | 4(50%) | 0.3(0.07-1.47) | 1.1(0.089-13.1) |
| 5. Single | 11(23.9%) | 35(76.1%) | Ref |  |
| 4. Occupational status |  |  |  |  |
| 1. House Servant | 1 (20%) | 4(80%) | 0.44(0.36-4.7) | 3.6(0.154-84.6) |
| 2. Daily Laborer | 14(11.6%) | 107(88.4%) | 0.8(0.51-8.3) | 2.1(0.51-8.3) |
| 3. Students | 5(41.7%) | 7(58.3%) | 0.16(0.04-0.65)* | 0.22(0.024-2) |
| 4. Farmer | 9(15%) | 51(85%) | 0.6(0.209-1.9) | 8.8(0.905-86.3) |
| 5. House wife | 4(13.3%) | 26(86.7%) | 0.7(0.187-2.8) | 1(0.14-7.4) |
| 6. Merchant | 6(9.4%) | 58(58%) | 0.7(0.187-2.8) | 1.7(0.36-8.11) |
| 7. Govt employee | 6(10%) | 54(90%) | Ref |  |

5. Educational status
|  |  |  |  |
| --- | --- | --- | --- |
| 1. Educated | 20(9.7%) | 186(90.3%) | 1.9(1.02-3.6)* |
| 2. Not educated | 25(17.1%) | 121(82.9%) | Ref |

\*significantly associated

The primary reasons reported by participants for treatment non-adherence were 15(33.3%) due to simply forgotten, 12(26.7%) being away from home, being busy 7(15. 6%).

Findings from in-depth interview also indicated that forgetfulness was mentioned by participants as barrier to adherence.

During the in-depth interview some patients forget their appointment dates and the dose of medication when they go to holy water and think of other spiritual and religious practice.

### Factors associated with medication adherence

The association of factors to medication adherence is reported in the tables (Table 3).Variables like age, gender, monthly income, active substance use, WHO clinical staging, CD4 count drug regimen and others variables were not associated with medication adherence.

Patients whose current viral load <1000 copies/μL were 11 times more (COR:10.9;:CI: 4.5- 26.6;P<0.0001), those who had side effects of ART drug 0.2 times less (COR: 0.193;CI: 0.01- 0.37;P<0.0001) likely were adherence to ART than their counter parts using bivariate analysis.

The association of health system related factor also indicated patients who came from nearby places (<5km or less) were 2.5 times more (COR: 2.5; CI: 1.3-5; P=0.006) in bivariate analysis. Selective variables were entered simultaneously into multivariate logistic regression to identify the most independent predictor variables of ART adherence.

Accordingly, the result showed that, married patients on ART were 3 times more (AOR: 3.2; CI: 1. 06-9.9;P=0.039) adherent than singles(Table,3).

Similarly, the in-depth interview indicated that, married people and patients who use memory aids were adherent due to the support and encouragement given from the spouse and family members.

Responses from the in-depth interviews with patients however indicated different findings. Some of them indicated that, alcohol consumption and being busy with social events were the reasons of forgetfulness to take their medication properly and influence adherence.

## Discussion

In this study we tried to assess the level of adherence and factors associated with HAART adherence in Debre Berhan Health Centre and Debre Berhan Referral Hospital, from February to April 2018.

Of the total respondents 87.2% of individuals has adherence level of > 95% to their ART drugs while 12.7% of them were non-adherent (<95%) and this finding is consistent with (61.8%,62.8%,75.4%,74.2%,63.8%,83.1%,86%,95.5%) studies done in Tawan Mai, an outpatient clinic, Taksin Hospital, Bangkok in Thailand, in Keffi, Nigeria, in Ibadan, Nigeria, Yirgalem hospital, Jimma University, in Bale Robe and Ambo (10-16). However it is lower than the finding (97.9%, 90%) of study conducted in South Eastern Nigeria, and Lagos Islan Nigeria (17, 18). And higher than (62.3%, 69%, 43.2%, 48.2%, 86%, 85.3%) Review study done in Africa 2014, studies done in Thailand (2010), Eldorate Kenya, Emu Kenya, Ghana and South Gondar ≥95% adherence rate respectively (18-24). This variation observed in different location may have different reasons, for example, It could be related to the heterogeneity in measurement methods since there is no consensus exists about the gold standard measurement of adherence Regarding psychosocial findings using bivariate analysis this study found that patients who disclosed their sero status to the family and community were 4.6 times higher likely adhered than not. On the other hand, patients who felt discomfort of taking medication in front of others were 0.5 times less likely to adhere than their counter parts in bivariate analysis. This finding is in line with the study conducted in Nigeria kefi, Ambo, Emmu Kenya and Bale Robe (11,15,16, 22).

In bivariate logistic regression analysis disclosure of HIV status to the family member shown 3.5 times more adherent than their counter parts similar other finding in which Respondent who disclosed their sero status to at least one person were 3.5 times more likely to be HAART adherent than those respondent who did not disclose their sero status (16). This is the same as finding in qualitative study. However, in the multivariate analysis no association was found which is similar with study in Mekele (25).

Using bivariate analysis and from in-depth interview Patients who got family support were more likely adhered than their counterparts which is consistent with studies in Diredawa and Harer (26). In this study the individuals who had support from the family was found to be 3.2 times likely to adhere in bivariate analysis. Similarly, findings in, Bale Robe, Gana, Jimma, North west Ethiopia and Diredawa also are in agreement with our finding (14,15, 24, 26). But in current study support was not significantly associated with adherence in the multivariate analysis and similar with study in Yirgalem, Bale robe, Gana and Diredawa (15,23,26, 27). This disappearance might be due to the effect of confounders.

This study also found that, patients who use memory aids like mobile, watch or other means of reminders were 6 times more likely to adherent than those who did not in bivariate and multivariate analysis but the odd of adherent in multivariate was decreased to 5. Our finding is not consistent with the finding from Nigeria which has shown the use of memory aids was not associated with medication adherence (11). But it was in agreement with the study that revealed the use of reminder tools are factors that influence adherence to ART and with current in-depth interview (28).

In this study current study there was association of current viral load with adherence, but no association was found between adherence and duration of infection, WHO clinical stage, base line CD4, current CD4 and baseline viral load. In contrary to this, baseline CD4 count of <350 cells/ml were significantly associated predictors of ART adherence as shown in Oromia (29). Longer time between HIV infection and AIDS had an important problem within the first six months of HAART adherence (30).

Generally, in the current study we identified that memory aids, marital status, side effect of medication and frequency of adherence to instruction were the main predictors of adherent to medication.

## Conclusions

Adherence level and certain important factors affecting adherence of patients have been explored and identified in this study.

The level of adherence to ART was relatively higher when compared to other studies done in Ethiopia and other developing countries but it is below the recommended level (≥95%) and multiple determinants of adherence were identified which needs to be addressed. The most frequently reported reason for non adherence were forgetting, went away from home, being busy with other things and do not want others to notice. The use of memory aids and being marred were found to be independent positive predictors for drug adherence whereas medication side effect and adherence instruction were the negative predictors for medication adherence. Additionally, ART dug stock out, long waiting time, use of alternative and traditional medicines, desire to live longer and fear of stigma and discrimination, were the findings reported as barriers to adherence.

## Acknowledgments

We are grateful to the Debre Berhan University for financial and logistic support. We would like to thank the hospital staff and administrators for their collaboration and unreserved help during data collection.

**Acronyms and List of abbreviations**
3TC: Lamivudine
ABC: Abacavir
AIDS: Acquired Immune Deficiency Syndrome
ART: Antiretroviral Therapy
D4T: Stavudine
DOT: Directly Observed Treatment
EFV: Efavirenz
LPV/r: Lopinavir/Ritonavir
MEMS: Medication Event Monitoring System
NNRTI: Non-Nucleoside Reverse-Transcriptase Inhibitor
NRTIs: Nucleoside Reverse Transcriptase Inhibitors
NVP: Neverapine
PI: Protease Inhibitors
PIT: Pill Identification Test
TDF: Tenofovir Disproxil Fumarate
TDM: Therapeutic Drug Monitoring
VAS: Visual Analogue Scale
ZDV: Zidovudine

## Ethics approval and consent to participate

Written consent was obtained from the Debre Berhan University (DBU) Health Science College and Debre Berhan town city admiration. Verbal consent obtained from each respondent. The confidentiality of the respondent respected throughout the procedures.

## Availability of data and material

The data set used for this study are available from the corresponding author at reasonable request.

## Competing interests

The authors declare that they have no competing interests.

## Funding

The author disclosed receipt of the following financial support for the research, authorship, and/or publication of this article: We are grateful to the Debre Berhan University for financial and logistic support.

## Authors’ contributions

All authors involved in preparing the proposal, analyses of the results and writing the final manuscript.

## References

1. Joint United Nation program on HIV/AIDS (UNAIDS). Prevention Gap Report. Geneva: unaids.org, 2016.

2. Joint United Nation Program on HIV/AIDS (USAIDS). Global Update on HIV Geneva: unaids.org, 2016.

3. World Health Organization(WHO). Global Update on HIV Treatment. Geneva: WHO, 2013.

4. Bottonari KA, Roberts JE, Ciesla JA, Hewitt RG. Life stress and adherence to antiretroviral therapy among HIV-positive individuals: a preliminary investigation. AIDS Patient Care STDS. 2005;19(11):719–27. Epub 2005/11/15.

5. World Health Organization(WHO). Guideline on When to Start Antiretroviral Therapy and on Pre-Exposure Prophylaxis For HiV. Geneva.: WHO; 2015.

6. Rubbert A, Behrens G, Ostrowsk M. Pathogenesis of HIV-1 infection in HIV Medicine. 2007:59–81.

7. World Health Organization (WHO). Global Update on HIV Evidence for action. Adherence to Long-Term Therapies.Geneva: WHO, 2003.

8. Steel G Nwokike J Mohan P. Development of a Multi-Method Tool to Measure ART Adherence in Resource-Constrained Settings: The South Africa Experience. United State of America (USA): Center for Pharmaceutical Management 2007:4–7.

9. World Health Organization (WHO). Antiretroviral Therapy for HIV Infection in Adults and Adolescents. Recommendations for a Public Health Approach. Austria: WHO; 2010.

10. Chamroonsawasdi K, Insri N, Pitikultang S. Predictive factors of antiretroviral (ARV) drug adherence among people living with HIV/AIDS attending at Taksin Hospital, Bangkok, Thailand. J Med Assoc Thai. 2011;94(7):775–81. Epub 2011/07/22.

11. Pennap GR, Abdullahi U, Bako IA. Adherence to highly active antiretroviral therapy and its challenges in people living with human immunodeficiency virus (HIV) infection in Keffi, Nigeria. JAHR. 2013;5(2):52–8,.

12. Omole Moses Kayode et al. Investigation of Factors Affecting Medication Adherence among People Living With Hiv/Aids under Non-Governmental Organizations in Ibadan City, Nigeria.JPBMS. 2012;21(21):1–5.

13. Markos E, Worku A, Davey G. Adherence to ART in PLWHA at Yirgalem Hospital, SouthEthiopia. EthiopJHealth Dev. 2008;22(2).

14. Abera A, Fenti B, Tesfaye T, Balcha F. Factors Influencing Adherence to Antiretroviral Therapy among People Living With HIV/AIDS at ART Clinic in Jimma University Teaching Hospital, Southwest Ethiopia. J Pharma 2015;1(1):2–6.

15. Mohammed AY, Ahmed MB, Tefera TB. Assessment of Factors Affecting Art Adherence among People Living with Human Immune Virus in Bale Robe Hospital, South East Ethiopia. ajphr. 2015;3(2):60–5.

16. Fituma S, Tsegaye D. Antiretroviral Therapy Adherence Among People Living With HIV In Ambo Hospital, West Shewa Zone, Oromia Region, Ethiopia. Obstet Gynecol Int J. 2016;Volume 5-(Issue 2):1–9

17. Ugochukwu U, Onyeonoro UU EU, Ibeh CC, Nwamoh UN, Ukegbu AU, Emelumadu OF,. Adherence to antiretroviral therapy among people living with human immunodeficiency virus/acquired immunodeficiency syndrome in a tertiary health facility in South Eastern Nigeria. J HIV Hum 2013;1(2):58–63.

18. Adekemi O, Sekoni, Obinna R, Obidik, Balogu M. Stigma, medication adherence and coping mechanism among people living with HIV attending General Hospital, Lagos Island, Nigeria. J Prm Health Care Fam. 2012:1–8.

19. Sung HK Sarah M Gerverb Sarah F Helen W. S.Adherence to Antiretroviral Therapy in Adolescents Living with HIV. Systematic Review and Meta-analysis:. AIDS. 2014;28(13): 20141–1950.

20. Li L, Lee SJ, Wen Y, Lin C, Wan D, Jiraphongsa C. Antiretroviral therapy adherence among patients living with HIV/AIDS in Thailand. Nurs Health Sci. 2010;12(2):212–20. Epub 2010/07/07.

21. Talam N Gatongi P Rotich J Kimaiyo S. Facctors Affecting Antiretroviral Drug Adherence among HIV/AIDS Adult Patients Attending HIV/AIDS Clinic at Moi Teaching and Referral Hospital, Eldoret, Kenya. East African Journal Of Public Health 2008:;5(2):1–77.

22. Kananu E, Mug N, Kabiru E, Mwaniki J. Patient Factors Influencing Adherence to ART Treatment among HIV/AIDS Patients in Embu Teaching and Referral Hospital Comprehensive Care Clinic. Science Journal of Public Health. 2016;4(5):75–380.

23. Obirikorang C, Selleh PK, Abledu JK, Fofie CO. Predictors of Adherence to Antiretroviral Therapy among HIV/AIDS Patients in the Upper West Region of Ghana. ISRN AIDS. 2013;2013:1–7.

24. Tadesse S, Tadesse A, Wubshet M. Adherence to Antiretroviral Treatment and Associated Factors among People Living with HIV/AIDS in Northwest Ethiopia. J Trop Dis. 2014;2(2):1–8.

25. Kiday Hailasillassiel, BE, Mussie, Alemayehul, Girmatsion, et al. Factors associated with adherence of highly active antiretro viral therapy among adult HIV/AIDS patients in Mekelle Hospital Northern Ethiopia. jsjph. 2014; 2(4):367–72.

26. Negesa L, Demeke E, Mekonnin W. Adherence to Antiretroviral Therapy and Factors affecting among People Living with HIV/AIDS and Taking Antiretroviral Therapy, Dire Dawa Town, Eastern Ethiopia. Journal of Infectious Diseases and Treatment.2017;3 (1: 5):1–6.

27. Markos E, Worku A, Davey G. Adherence to ART in PLWHA at Yirgalem Hospital, SouthEthiopia. EthiopJHealth Dev. 2008;22(2).

28. Chen Y, Kalichman SC S. Synergistic effects of food insecurity and drug use on medication adherence among people living with HIV infection. J Behav Med. 2015;38:397–406.

29. Chaka TE, Abeya SG, Adlo AM, Abebe TW, Hamuse SD, Lencha MT, et al. Antiretroviral Therapy: Level of Adherence and Its Determinants Among Patients on Treatment in Different Health Facilities. A Cross Sectional Study in Oromia Regional State, Ethiopia. J AIDS Clin Res. 2016;7(11):1–7.

30. Silva JAG, Dourad I, Brito AMd, Silva CALd. Factors associated with non-adherence to antiretroviral therapy in adults with AIDS in the first six months of treatment in Salvador, Bahia State, Brazil. Rio de Janeiro. 2015;31(6):1–11.

